# Differences between intrinsic and acquired nucleoside analogue resistance in acute myeloid leukaemia cells

**DOI:** 10.1101/2021.07.19.452885

**Authors:** Tamara Rothenburger, Dominique Thomas, Yannick Schreiber, Paul R. Wratil, Tamara Pflantz, Kirsten Knecht, Katie Digianantonio, Joshua Temple, Constanze Schneider, Hanna-Mari Baldauf, Katie-May McLaughlin, Florian Rothweiler, Berna Bilen, Samira Farmand, Denisa Bojkova, Rui Costa, Nerea Ferreirós, Gerd Geisslinger, Thomas Oellerich, Yong Xiong, Oliver T. Keppler, Mark N. Wass, Martin Michaelis, Jindrich Cinatl

## Abstract

**Background:** SAMHD1 mediates resistance to anti-cancer nucleoside analogues, including cytarabine, decitabine, and nelarabine that are commonly used for the treatment of leukaemia, through cleavage of their triphosphorylated forms. Hence, SAMHD1 inhibitors are promising candidates for the sensitisation of leukaemia cells to nucleoside analogue-based therapy. Here, we investigated the effects of the cytosine analogue CNDAC, which has been proposed to be a SAMHD1 substrate, in the context of SAMHD1.

**Methods:** CNDAC was tested in 13 acute myeloid leukaemia (AML cell lines), in 26 acute lymphoblastic leukaemia cell lines, ten AML sublines adapted to various antileukaemic drugs, 24 single cell-derived clonal AML sublines, and primary leukaemic blasts from 24 AML patients. Moreover, 24 CNDAC-resistant sublines of the AML cell lines HL-60 and PL-21 were established. The *SAMHD1* gene was disrupted using CRISPR/Cas9 and SAMHD1 depleted using RNAi, and the viral Vpx protein. Forced DCK expression was achieved by lentiviral transduction. SAMHD1 promoter methylation was determined by PCR after treatment of genomic DNA with the methylation-sensitive HpaII endonuclease. Nucleoside (analogue) triphosphate levels were determined by LC-MS/MS. CNDAC interaction with SAMHD1 was analysed by an enzymatic assay and by crystallisation.

**Results:** Although the cytosine analogue CNDAC was anticipated to inhibit SAMHD1, SAMHD1 mediated intrinsic CNDAC resistance in leukaemia cells. Accordingly, SAMHD1 depletion increased CNDAC triphosphate (CNDAC-TP) levels and CNDAC toxicity. Enzymatic assays and crystallisation studies confirmed CNDAC-TP to be a SAMHD1 substrate. In 24 CNDAC-adapted acute myeloid leukaemia (AML) sublines, resistance was driven by DCK (catalyses initial nucleoside phosphorylation) loss. CNDAC-adapted sublines displayed cross-resistance only to other DCK substrates (e.g. cytarabine, decitabine). Cell lines adapted to drugs not affected by DCK or SAMHD1 remained CNDAC sensitive. In cytarabine-adapted AML cells, increased SAMHD1 and reduced DCK levels contributed to cytarabine and CNDAC resistance.

**Conclusion:** Intrinsic and acquired resistance to CNDAC and related nucleoside analogues are driven by different mechanisms. The lack of cross-resistance between SAMHD1/ DCK substrates and non-substrates provides scope for next-line therapies after treatment failure.

## Introduction

Drug resistance is a main obstacle in the successful treatment of cancer [Fenton et al., 2018; Michaelis et al., 2019; Bukowski et al., 2020]. Resistance can be either intrinsic or acquired. Intrinsic resistance means that a therapy-naïve cancer does not respond to treatment right from the start. In acquired resistance, there is an initial therapy response, but resistance develops over time [Michaelis et al., 2019; Santoni-Rugiu et al., 2019].

Intrinsic and acquired resistance are conceptually different. Intrinsic resistance is a collateral event during carcinogenesis not influenced by treatment. In contrast, acquired resistance is the consequence of a directed evolution driven by therapy. In agreement, discrepancies have been detected between drug resistance mechanisms in the intrinsic and the acquired resistance setting [Michaelis et al., 2019; Oellerich et al., 2019; Santoni-Rugiu et al., 2019; Touat et al., 2020].

Sterile alpha motif and histidine-aspartate domain-containing protein 1 (SAMHD1) is a deoxynucleoside triphosphate (dNTP) triphosphohydrolase that cleaves physiological dNTPs into deoxyribonucleotides and inorganic triphosphate [Goldstone et al., 2011; Powell et al., 2011]. SAMHD1 also inactivates the triphosphorylated forms of some anti-cancer nucleoside analogues [Schneider et al., 2017; Herold et al., 2017; Knecht et al., 2018; Oellerich et al., 2019; Rothenburger et al., 2020; Xagoraris et al., 2021]. High SAMHD1 levels indicate poor clinical response to nucleoside analogues such as cytarabine, decitabine, and nelarabine in acute myeloid leukaemia (AML), acute lymphoblastic leukaemia, and Hodgkin lymphoma [Schneider et al., 2017; Oellerich et al., 2019; Rothenburger et al., 2020; Xagoraris et al., 2021]. Moreover, previous findings indicated differing roles of SAMHD1 in intrinsic and acquired resistance to nucleoside analogues [Schneider et al., 2017; Oellerich et al., 2019].

Here, we investigated intrinsic and acquired resistance against the nucleoside analogue 2′-C-cyano-2′-deoxy-1-β-*D*-arabino-pentofuranosyl-cytosine (CNDAC). CNDAC and its orally available prodrug sapacitabine display clinical activity against AML [Kantarjian et al., 2010; Kantarjian et al., 2012; Czmerska et al., 2018; Kantarjian et al., 2019]. We selected CNDAC, because, in contrast to SAMHD1 substrates such as cytarabine and decitabine, it has been proposed to be a SAMHD1 inhibitor [Hollenbaugh et al., 2017]. CNDAC is further interesting due to its unique mechanism of action among deoxycytidine analogues, which is characterised by CNDAC triphosphate (CNDAC-TP) incorporation into DNA initially causing single strand breaks and G2 cell cycle arrest [Hanaoka et al., 1999; Azuma et al., 2001; Liu et al., 2005; Liu et al., 2008; Al Abo et al., 2017; Liu et al., 2018; Liu et al., 2019].

## Results

### SAMHD1 levels correlate with leukaemia cell sensitivity to CNDAC

Initially, we characterised a panel of 13 human AML cell lines for the levels of SAMHD1 and deoxycytidine kinase (DCK) (Figure 1A). DCK phosphorylates and activates cytidine analogues in a rate-limiting step [Lotfi et al., 2003; Homminga et al., 2011; Wu et al., 2021] and may, hence, determine cell sensitivity to a nucleoside analogue like CNDAC anticipated to be a SAMHD1 inhibitor [Hollenbaugh et al., 2017]. We detected varying SAMHD1 and DCK levels (Figure 1A, Suppl. Figure 1), varying CNDAC concentrations that reduced cell viability by 50% (IC_50_) (Figure 1B, Suppl. Figure 2, Suppl. Table 1), and varying CNDAC-TP levels (Figure 1C) across the investigated cell lines. However, the CNDAC IC_50_s did not correlate with the cellular levels of DCK (Figure 1D), indicating that DCK is not a critical determinant of CNDAC activity in our cell line panel.

**Figure 1.**
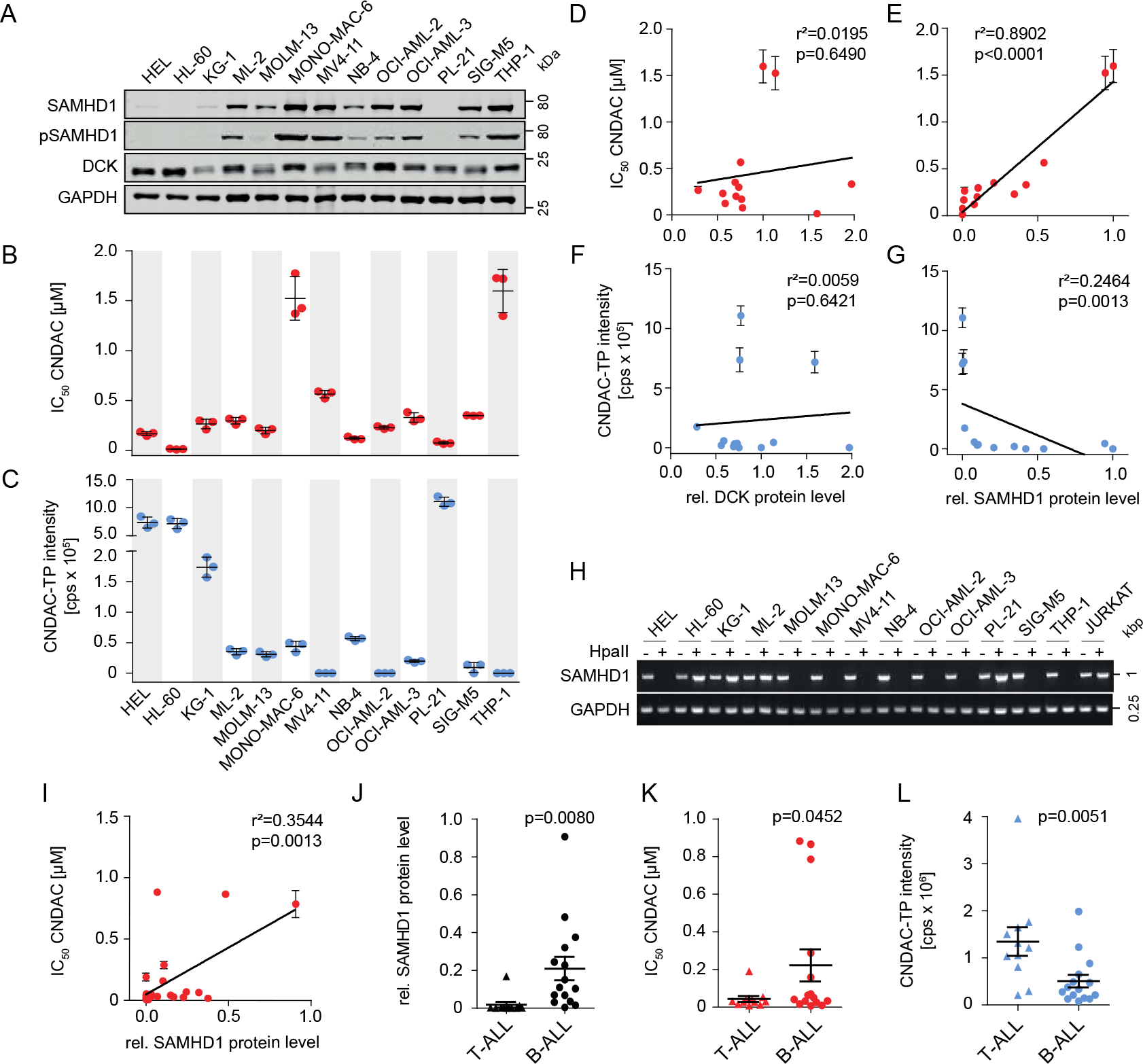
SAMHD1 (but not DCK) levels determine sensitivity to CNDAC and inversely correlate with CNDAC-triphosphate (CNDAC-TP) in leukaemia cell lines. (A) Representative Western blots of SAMHD1, phosphorylated SAMHD1 (pSAMHD1), and DCK in 13 AML cell lines. GAPDH served as loading control. Uncropped Western blots are presented in Supplementary Figure 1. (B) CNDAC concentrations that reduce the viability of AML cell lines by 50% (IC_50_). Horizontal lines and error bars represent means ± SD of three independent experiments. (C,D) Correlation of the CNDAC IC_50_ values with cellular DCK (C) or SAMHD1 (D) protein levels, quantified using near-infrared Western blot images to determine the ratio DCK/ GAPDH or SAMHD1/ GAPDH. Closed circles and error bars represent means ± SD of three independent experiments. Linear regression analyses were performed using GraphPad Prism. (E) CNDAC triphosphate (CNDAC-TP) levels determined by LC–MS/MS. Horizontal lines and error bars show means ± SD of three independent experiments. (F,G) Correlation of CNDAC-TP levels with cellular DCK (F) or SAMHD1 (G) protein levels in AML cell lines, quantified using near-infrared Western blot images to determine the ratio DCK/ GAPDH or SAMHD1/ GAPDH. Closed circles and error bars represent means ± SD of three independent experiments. Linear regression analyses were performed using GraphPad Prism. (H) Analysis of *SAMHD1* promoter methylation in AML cell lines through amplification of a single PCR product (993-bp) corresponding to the promoter sequence after *Hpa*II digestion. A 0.25-kb fragment of the GAPDH gene lacking *Hpa*II sites was PCR-amplified using the same template DNA served as loading control. THP-1 served as control cell for an unmethylated *SAMHD1* promotor, while JURKAT served as control cell for a methylated promotor. (I) Correlation of CNDAC IC_50_ values in 26 ALL cell lines (11 T-ALL, 15 B-ALL) with SAMHD1 protein levels, quantified using near-infrared Western blot images to determine the ratio SAMHD1/ GAPDH relative to the positive control THP-1. Closed circles and error bars represent means ± SD of three independent experiments. Linear regression analyses were performed using GraphPad Prism. (J-L) Comparison of SAMHD1 protein levels (J), CNDAC IC_50_ values (K) and CNDAC-TP levels determined by LC-MS/MS (L) in T-ALL and B-ALL cells. Each point represents the mean of three independent experiments. One-tailed Student’s t-tests were used to compare means in T-ALL and B-ALL cells (represented as horizontal lines ± SEM).

In contrast, the CNDAC IC_50_s correlated with the cellular SAMHD1 levels (Figure 1E), suggesting that SAMHD1 may cleave and inactivate CNDAC-TP, although CNDAC had been proposed to be a SAMHD1 inhibitor [Hollenbaugh et al., 2017]. Also, there was no correlation between cellular CNDAC-TP and DCK levels (Figure 1F), but an inverse correlation of the CNDAC-TP levels with SAMHD1 (Figure 1G). This further supports the notion that SAMHD1 but not DCK critically determines CNDAC phosphorylation and activity. Notably, SAMHD1 promoter methylation (Figure 1H) did not always indicate cellular SAMHD1 levels (Figure 1A), showing that multiple mechanisms are involved in regulating the cellular abundance of this protein.

The CNDAC IC_50_s also correlated with the cellular SAMHD1 levels in acute lymphoblastic leukaemia (ALL) cells (Figure 1I, Suppl. Table 2). In agreement with previous findings [Rothenburger et al. 2020], T-cell ALL (T-ALL) cells were characterised by lower SAMHD1 levels than B-ALL cells (Figure 1J). This was reflected by higher CNDAC sensitivity (Figure 1K) and higher CNDAC-TP levels (Figure 1L) in T-ALL cells than in B-ALL cells. Taken together, these findings suggest that CNDAC is a SAMHD1 substrate and that SAMHD1 but not DCK critically determines CNDAC phosphorylation and activity in AML and ALL cells.

### SAMHD1 suppression sensitises leukaemia cells to CNDAC

Functional studies further confirmed the impact of SAMHD1 on CNDAC activity. THP-1 AML cells, in which the *SAMHD1* gene was disrupted using CRISPR/Cas9 (THP-1 KO cells), displayed increased CNDAC sensitivity (Figure 2A) and CNDAC-TP levels (Figure 2B) relative to control cells. Moreover, THP-1 KO cells showed enhanced DNA damage, as indicated by γH2AX levels, CHK2 phosphorylation, and TIF1β phosphorylation (Figure 2C), and apoptosis, as indicated by PARP cleavage (Figure 2C) and caspase 3/7 activity (Figure 2D, Suppl. Table 3), in response to CNDAC. This is in line with the anticipated mechanism of action of CNDAC, i.e. CNDAC-TP incorporation into DNA resulting in strand breaks and apoptosis [Hanaoka et al., 1999; Azuma et al., 2001; Liu et al., 2005; Liu et al., 2008; Al Abo et al., 2017; Liu et al., 2018; Liu et al., 2019].

**Figure 2.**
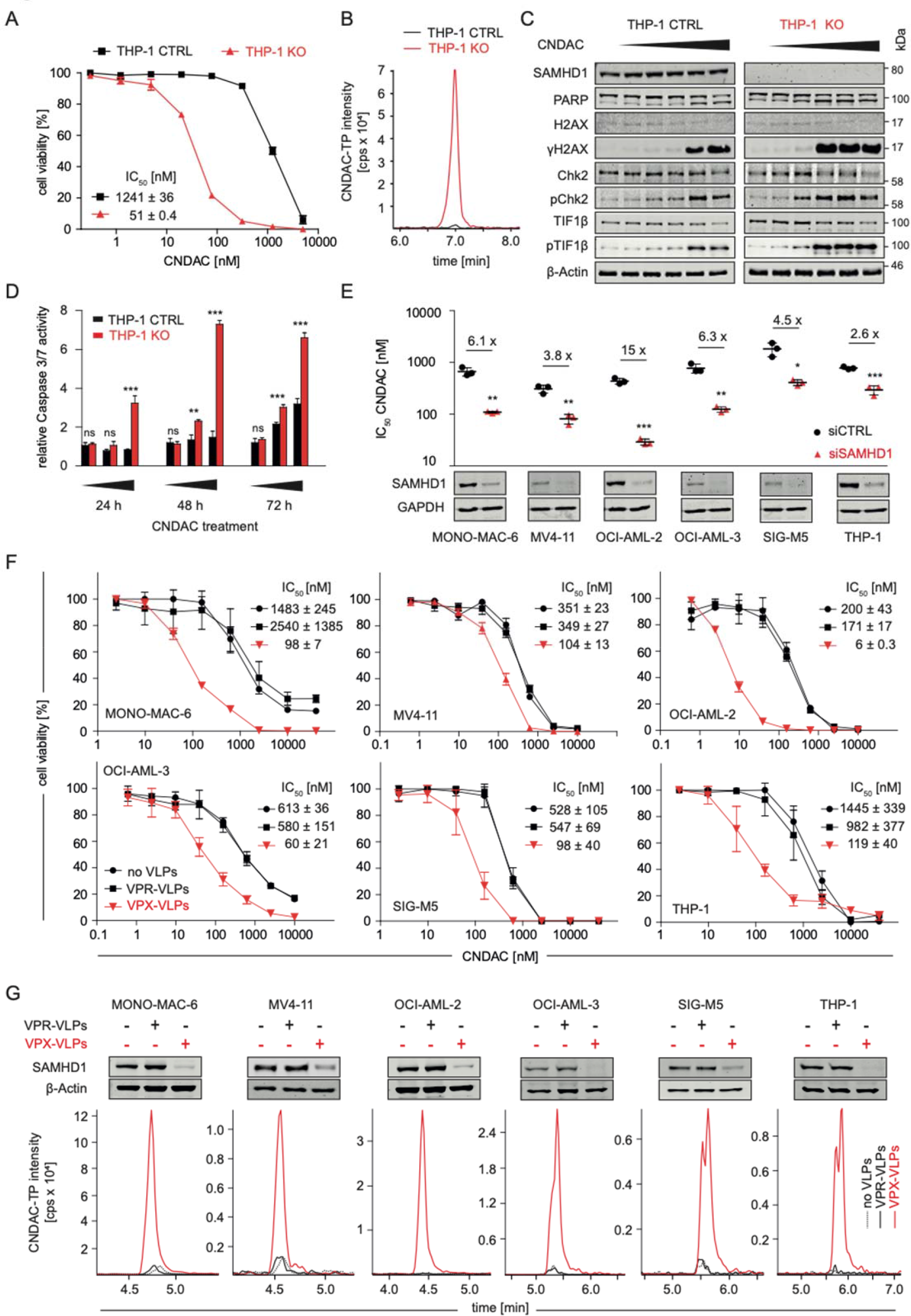
SAMHD1 suppression sensitises AML cells to CNDAC. (A) CNDAC dose-response curves in THP-1 knockout (THP-1 KO) cells, in which the *THP-1* gene was disrupted using CRISPR–Cas9, or control cells (THP-1 CTRL). Values represent means ± SD of three independent experiments. Concentrations that reduce cell viability by 50% (IC_50_s) ± SD are provided. (B) Representative LC-MS/MS analysis of CNDAC triphosphate (CNDAC-TP) levels in THP-1 KO and THP-1 CTRL cells. (C) Representative Western blots indicating levels of proteins involved in DNA damage response in THP-1 KO and THP-1 CTRL cells after treatment with increasing CNDAC concentrations (0, 3.2, 16, 80, 400, and 2000 nM) for 72 hours. (D) Caspase 3/7 activity in THP-1 KO and THP-1 CTRL cells after treatment with increasing CNDAC concentrations (0.015, 0.9375 and 60 µM) for 24, 48, and 72 hours, relative to untreated controls. Mean ± SD is provided for one representative experiment out of three using three technical replicates. p-values were determined by two-tailed Student’s t-tests (*p < 0.05; **p < 0.01; ***p < 0.001). (E) CNDAC IC_50_ values in AML cells after transfection with SAMHD1-siRNAs (siSAMHD1) or non-targeting control siRNAs (siCTRL). Values represent the means ± SD of three technical replicates of one representative experiment out of three. p-values were determined by two-tailed Student’s t-tests (*p < 0.05; **p < 0.01; ***p < 0.001). (F) CNDAC dose-response curves in AML cell lines treated with CNDAC in the absence or presence of VPX virus-like particles (VPX-VLPs, cause SAMHD1 depletion), or VPR virus-like particles (VPR-VLPs, negative control). Values represent the means ± SD of three technical replicates of one representative experiment out of three. (G) Representative Western Blots and LC-MS/MS analyses of CNDAC-TP levels in AML cells treated with VPX-VLPs or control VPR-VLPs.

Further, SAMHD1 depletion using siRNA (Figure 2E, Suppl. Figure 3) and virus-like particles (VLPs) carrying the lentiviral VPX protein (VPX-VLPs) [Schneider et al., 2017] (Figure 2F) increased the CNDAC sensitivity of AML cell lines. VPX-VLP-mediated SAMHD1 depletion was also associated with elevated CNDAC-TP levels (Figure 2G). These findings further support a critical role of SAMHD1 in determining CNDAC sensitivity of AML cells.

### SAMHD1 determines sensitivity of primary AML cells

CNDAC sensitivity also correlated with the cellular SAMHD1 levels in primary leukaemic blasts derived from the bone marrow of 24 therapy-naïve AML patients (Figure 3A, Suppl. Figure 4, Suppl. Table 4). Moreover, primary leukaemic blasts were sensitised by VPX-VLPs to CNDAC (Figure 3B, Figure 3C, Suppl. Figure 5) and VPX-VLP-mediated SAMHD1 depletion resulted in increased CNDAC-TP levels in AML blasts (Figure 3D, Figure 3E). This shows that SAMHD1 also determines CNDAC sensitivity in clinical AML samples.

**Figure 3.**
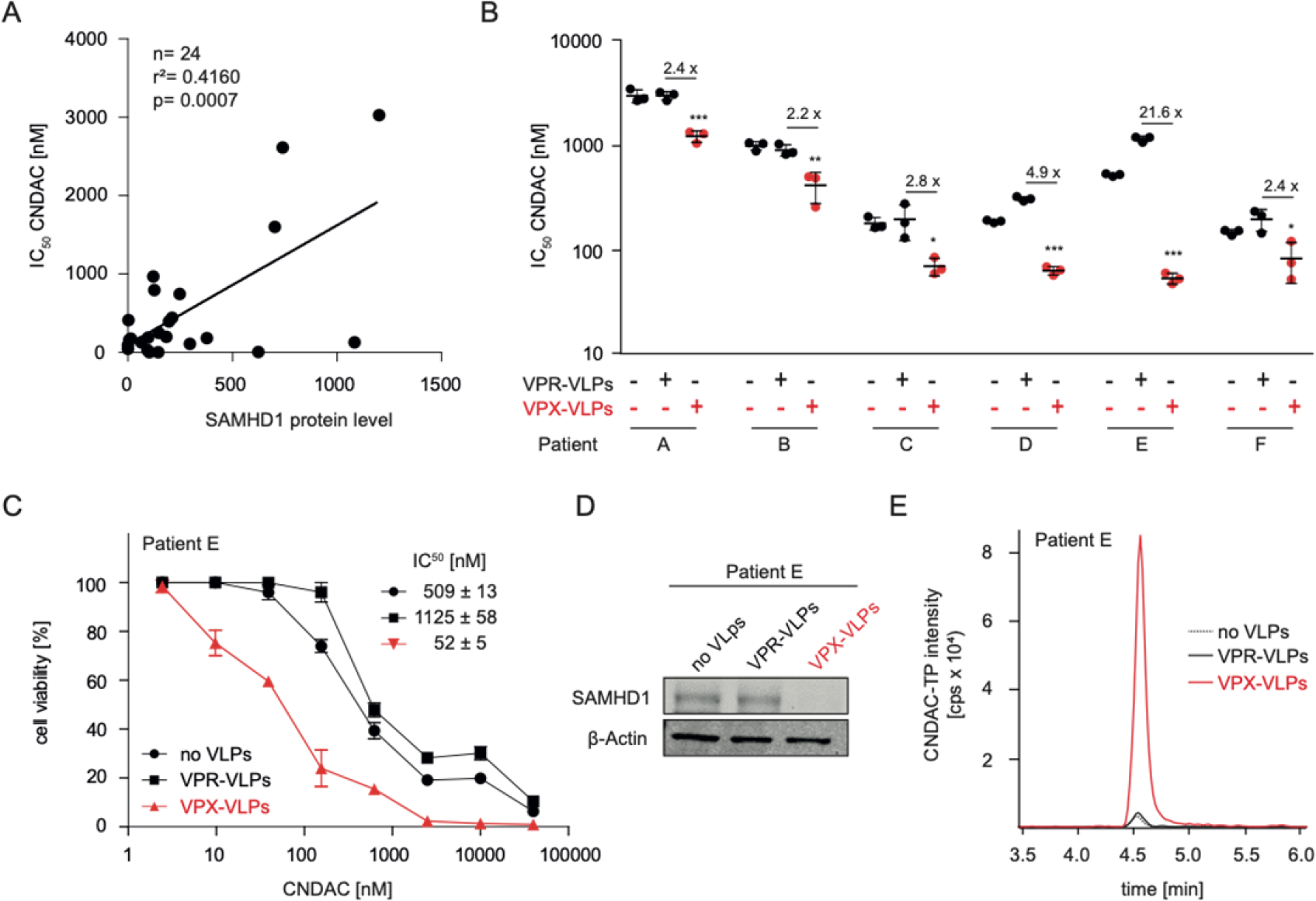
SAMHD1 determines CNDAC sensitivity of primary AML cells. (A) Correlation of SAMHD1 protein levels and CNDAC concentrations that reduce cell viability by 50% (IC_50_s) in bone-marrow-derived leukaemic blasts derived from 24 therapy-naïve AML patients. Cells were co-immunostained for CD33, CD34, CD45 (surface markers) and intracellular SAMHD1 and the mean fluorescence intensity (MFI) was analysed by flow cytometry. ATP assays were performed in three technical replicates to determine the CNDAC IC50 values. Linear regression analyses were performed using GraphPad Prism. (B) CNDAC IC_50_ values in bone-marrow-derived leukaemic blasts derived from six therapy-naïve AML patients either treated with VPX virus-like particles (VPX-VLPs, cause SAMHD1 depletion), VPR virus-like particles (VPR-VLPs, negative control), or left untreated. Horizontal lines and error bars indicate means ± SD of three technical replicates. p-values were determined by two-tailed Student’s t-tests (*p < 0.05; **p < 0.01; ***p < 0.001). (C) CNDAC dose-response curves in primary AML cells of one exemplary patient (Patient E) treated with VPX-VLPs, VPR-VLPs or left untreated. IC_50_ values represent means ± SD of three technical replicates. (D) Representative Western blots indicating SAMHD1 levels in primary AML cells derived from Patient E in response to treatment with VPX-VLPs. (E) CNDAC-triphosphate (CNDAC-TP) levels as determined by LC-MS/MS in primary AML cells derived from Patient E in response to treatment with VPX-VLPs.

### SAMHD1 hydrolyses CNDAC triphosphate (CNDAC-TP)

Next, we studied the interaction of CNDAC-TP and SAMHD1 in an enzymatic assay. SAMHD1 forms a homotetramer complex that cleaves nucleoside triphosphate (Suppl. Figure 6). Tetramer formation depends on nucleoside triphosphate binding to the allosteric SAMHD1 sites 1 (A1) and A2. A1 is activated by guanosine triphosphate (GTP) or desoxy-guanosine triphosphate (dGTP) binding. A2 can be activated by any canonical deoxy-nucleoside triphospate (dNTP) and some triphosphorylated deoxyribose-based nucleoside analogues such as cladribine-TP and decitabine-TP (Suppl. Figure 6) [Ji et al., 2013; Ji et al., 2014; Hollenbaugh et al., 2017; Knecht et al., 2018; Oellerich et al., 2019]. Arabinose-based nucleoside analogue triphosphates (e.g. cytarabine-TP, fludarabine-TP, or arabinosylguanine-TP (AraG-TP, the active metabolite of nelarabine), and the triphosphorylated 2’-deoxy-2’-fluororibose-based nucleoside analogue clofarabine depend on the activation of A2 by canonical nucleotides [Hollenbaugh et al., 2017; Knecht et al., 2018; Oellerich et al., 2019].

Results from the enzymatic assay confirmed that SAMHD1 hydrolyses CNDAC-TP only in the presence of dGTP (Figure 4A). This indicates that CNDAC-TP is a SAMHD1 substrate but not able to activate the enzyme by binding to A1 and A2.

**Figure 4.**
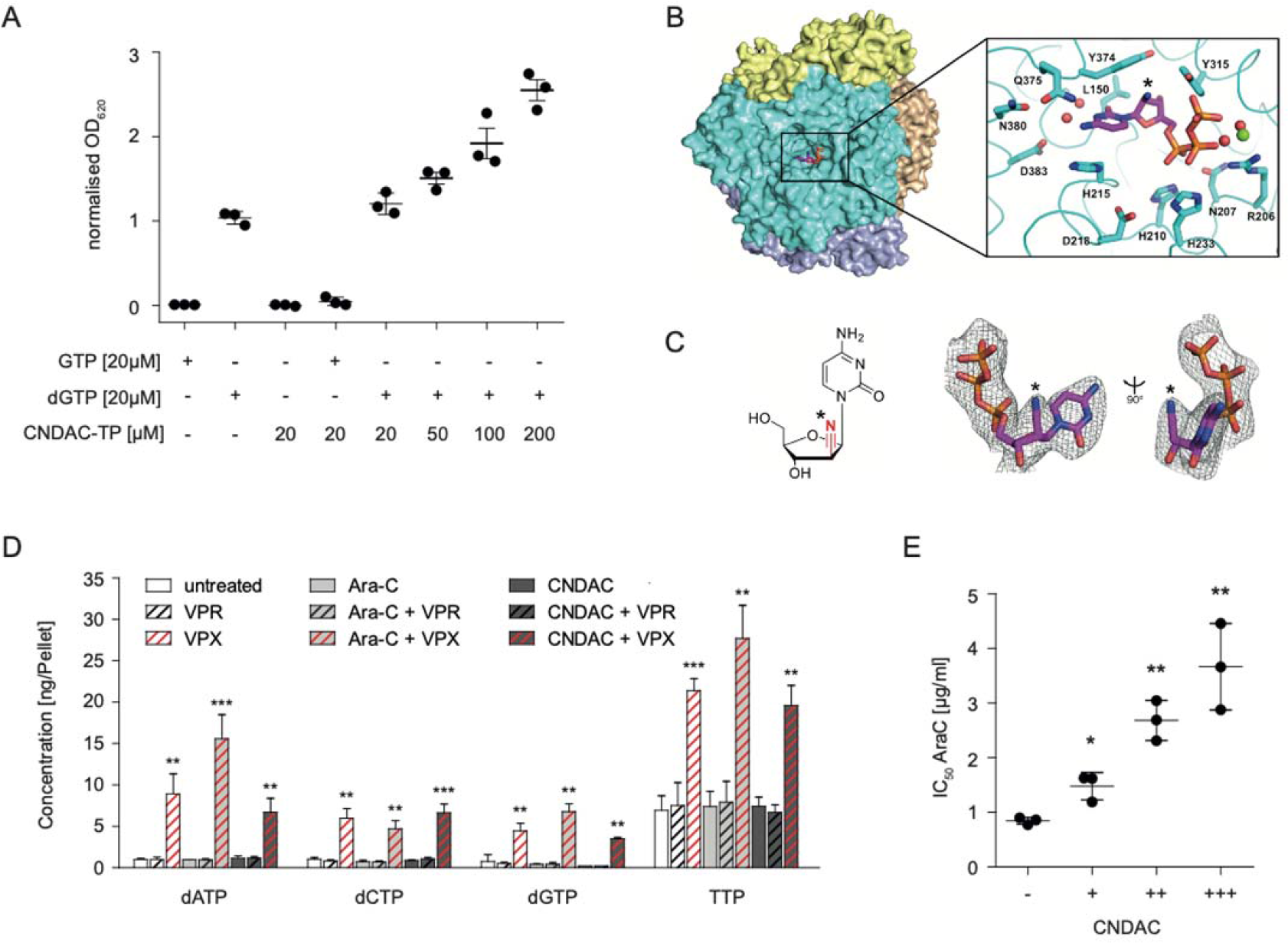
CNDAC triphosphate (CNDAC-TP) is a SAMHD1 substrate. (A) Normalised results from a colorimetric SAMHD1 activity assay carried out in presence of different combinations of GTP, dGTP and CNDAC-TP. Horizontal lines and error bars represent means ± SD from three independent experiments. (B) Surface view of SAMHD1 tetramer with each subunit in a different colour. CNDAC-TP in a catalytic pocket is shown in magenta sticks. (B, inset) CNDAC-TP bound to the SAMHD1 catalytic pocket. Black asterisks indicate the site of nitrile modification. The SAMHD1 backbone is shown as ribbons with side chains shown as sticks. A magnesium ion is shown as a green sphere and coordinated waters are shown as red spheres. Portions of the structure are omitted for clarity. (C) Chemical structure of CNDAC with 2’S nitrile modification highlighted (left). 2Fo-Fc electron density (σ = 1.0) for CNDAC-TP co-crystallized in the catalytic pocket of SAMHD1 (right). Black asterisks indicate site of nitrile modification. (D) Concentrations of physiological dNTPs in THP-1 cells determined by LC-MS/MS after pre-treatment with VPX virus-like particles (VPX-VLPs, cause SAMHD1 depletion), VPR virus-like particles (VPR-VLPs, negative control), and with or without cytarabine (AraC) or CNDACs. Bars and error bars represent means ± SD from three independent measurements. The Lower Limit of Quantification (LLOQ) for dGTP was 0.2 ng/Pellet, so values below the LLOQ were set to 0.2 ng/Pellet (CNDAC and CNDAC + VPR). p-values were determined by two-tailed Student’s t-tests were performed (*p < 0.05; **p < 0.01; ***p < 0.001). (E) AraC IC_50_s in THP-1 cells in the presence of different CNDAC concentrations (0, 0.375, 0.75, 1.5 µM). Horizontal lines and error bars represent means ± SD of three technical replicates of one representative experiment out of three. p-values were determined by two-tailed Student’s t-tests were performed (*p < 0.05; **p < 0.01; ***p < 0.001).

### Crystal structure of CNDAC-TP bound to SAMHD1

To investigate the interaction of CNDAC-TP and SAMHD1 further, we crystallised the catalytically inactive HD domain (residues 113-626; H206R, D207N) of SAMHD1 in the presence of GTP, dATP, and excess CNDAC-TP as previously described [Knecht et al., 2018] and collected diffraction data to 2.8Å. SAMHD1 crystallised as a tetramer with GTP and dATP occupying A1 and A2, respectively, and CNDAC-TP bound to the catalytic site (Figure 4B, Suppl. Table 5).

Previous studies investigating the binding of triphosphorylated nucleoside analogues to SAMHD1 showed that modifications at the 2’ribose (R) position are major determinants of interaction with the catalytic SAMHD1 site [Knecht et al, 2018]. Analogues with 2’R modifications abrogate binding to SAMHD1, while 2’S stereoisomers are more permissive. Furthermore, the catalytic site tolerates larger 2’S modifications, whereas analogue binding at the A2 site is either impaired or fully blocked by 2’S fluorination or hydroxylation of the sugar ring, respectively [Knecht et al., 2018].

Consistent with these observations, the CNDAC-TP-bound SAMHD1 adopts the same conformation as the canonical nucleotide-bound form (overall RMSD: 0.30 Å vs PDB ID 4BZB). The ribose 2’S nitrile modification of CNDAC-TP (Figure 4C) protrudes outward from the catalytic pocket without affecting canonical nucleotide contacts with active site residues. CNDAC-TP is therefore easily accommodated in the catalytic site to serve as a substrate for SAMHD1 triphosphohydrolase activity. However, the large nitrile group of CNDAC-TP prevents binding to the more restrictive A2 site. Thus, CNDAC alone is insufficient for SAMHD1 activation.

### Impact of CNDAC on cellular levels of physiological nucleoside triphosphates and the activity of SAMHD1 substrates

The finding that CNDAC-TP is itself a substrate of SAMHD1 does not exclude the possibility that it also exerts inhibitory effects on SAMHD1, as previously suggested [Hollenbaugh et al., 2017]. Hence, we investigated the effects of CNDAC on the levels of physiological desoxynucleoside triphosphates (dNTPs) and the activity of cytarabine, the triphosphate of which is known to be a SAMHD1 substrate [Schneider et al., 2017].

CNDAC did in contrast to VPX-VLPs, which served as a positive control for suppressing SAMHD1 activity, not increase the levels of physiological dNTPs (Figure 4D). Moreover, CNDAC did not increase the activity of cytarabine (Figure 4E). Thus, these findings do not suggest a pharmacologically relevant activity of CNDAC as SAMHD1 inhibitor in AML cells.

### Clonal heterogeneity in SAMHD1 levels drives intrinsic AML cell resistance to CNDAC

When we established twelve single cell-derived clones of the AML cell line MV4-11 by limited dilution (Figure 5A), we determined an up to 332-fold difference in CNDAC sensitivity (CNDAC IC_50_ clone 1: 0.065 µM, CNDAC IC_50_ clone 11: 21.6 µM; Figure 5B, Suppl. Figure 7). Moreover, the MV4-11 clones displayed substantial discrepancies in the cellular SAMHD1 levels (Figure 5C), but no changes in SAMHD1 promoter methylation (Figure 5D).

**Figure 5.**
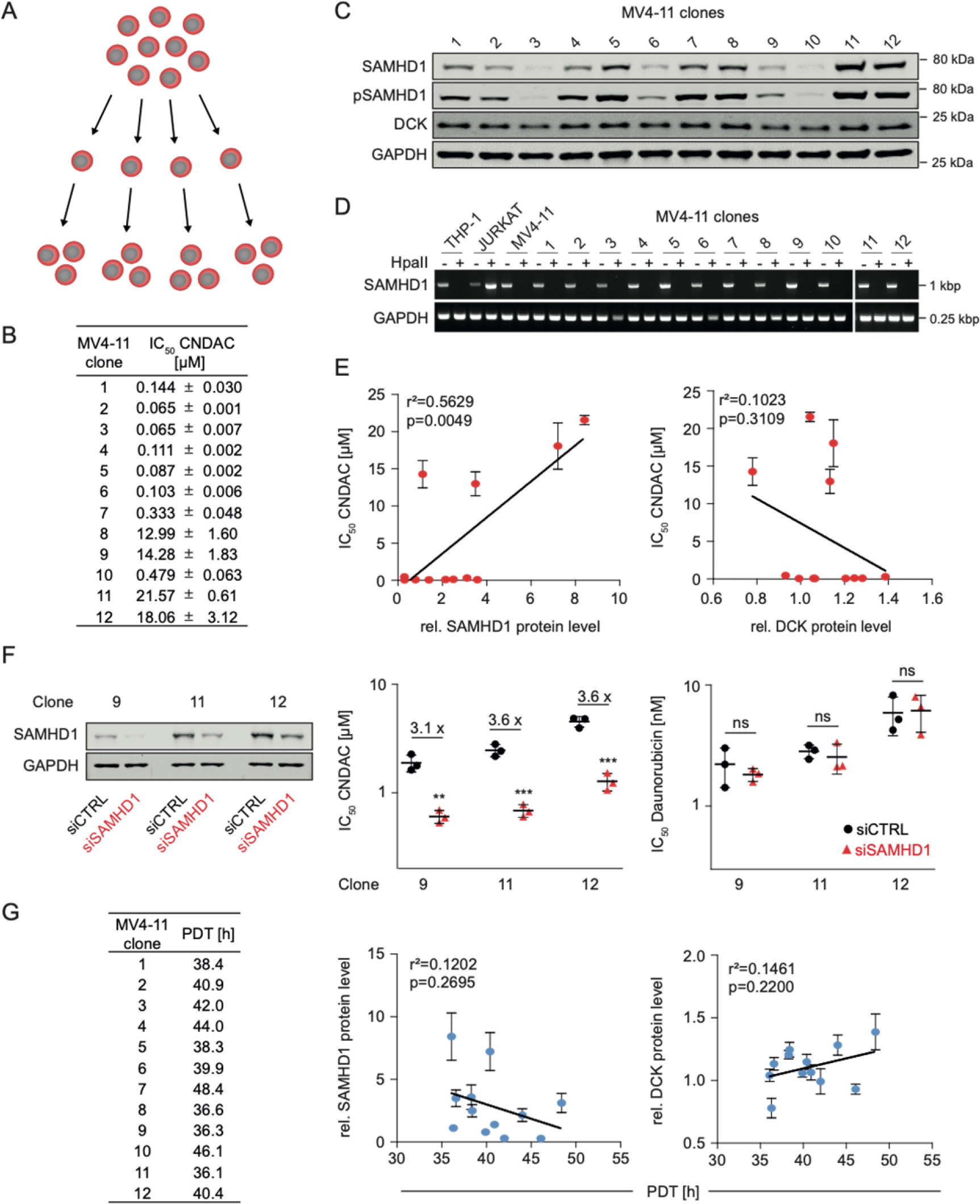
Clonal heterogeneity in SAMHD1 levels drives intrinsic resistance to CNDAC but not population doubling time in MV4-11 cells. (A) Schematic illustration of the establishment of MV4-11 single cell-derived clones by limited dilution. (B) CNDAC concentrations that reduce viability of 12 single-cell-derived MV4-11 clones by 50% (IC_50_). Values represent means ± SD of three independent experiments. (C) Representative Western blots of SAMHD1, phosphorylated SAMHD1 (pSAMHD1), and DCK in single cell-derived MV4-11 clones. GAPDH served as a loading control. (D) Analysis of *SAMHD1* promoter methylation in MV4-11 clones through amplification of a single PCR product (993-bp) corresponding to the promoter sequence after *Hpa*II digestion. (E) Correlation of the CNDAC IC_50_ values with cellular SAMHD1 or DCK protein levels, quantified using near-infrared Western blot images to determine the ratio SAMHD1/ GAPDH or DCK/ GAPDH. Closed circles and error bars represent means ± SD of three independent experiments, each performed in three technical replicates. Linear regression analyses were performed using GraphPad Prism. (F) Western Blots and IC_50_ values for CNDAC and Daunorubicin in MV4-11 clones 9, 11, and 12 after transfection with SAMHD1-siRNAs (siSAMHD1) or non-targeting control siRNAs (siCTRL). Each symbol represents the mean ± SD of three technical replicates of one representative experiment out of three. P-values were determined by two-tailed Student’s t-test (*p < 0.05; **p < 0.01; ***p < 0.001). (G) Population doubling time (PDT) in MV4-11 single cell-derived clones and correlation of the PDT with cellular SAMHD1 or DCK protein levels. Closed circles and error bars represent means ± SD from the quantification of three Western Blots. Linear regression analyses were performed using GraphPad Prism.

There was a significant correlation between SAMHD1 protein levels (but not the DCK protein levels) and the CNDAC IC_50_s (Figure 5E), and siRNA-mediated SAMHD1 depletion resulted in increased CNDAC (but not daunorubicin) sensitivity in three selected clones displaying differing SAMHD1 levels (Figure 5F, Suppl. Figure 8). The different effects of SAMHD1 on CNDAC- and daunorubicin-mediated toxicity suggest that SAMHD1 interferes with CNDAC activity predominantly by cleaving CNDAC-TP and not by generally augmenting DNA repair.

Differences in cellular SAMHD1 levels may affect cell proliferation [Franzolin et al., 2013; Kohnken et al., 2015; Kodigepalli et al., 2018; Wu Y et al., 2021], but there was no significant correlation between the SAMHD1 (or DCK) levels of the MV4-11 clones and their doubling times (Figure 5G).

Taken together, these findings confirm that the response to CNDAC is primarily driven by the SAMHD1 levels in CNDAC-naïve AML cells.

### Acquired resistance to CNDAC is associated with decreased DCK levels

To investigate the role of SAMHD1 in acquired CNDAC resistance, we established twelve CNDAC-resistant sublines of each of the AML cell lines HL60 and PL21, which are characterised by low SAMHD1 levels (Figure 1A) and high CNDAC sensitivity (Figure 1B). Interestingly, none of the 24 resulting CNDAC-resistant sublines displayed increased SAMHD1 levels but all showed reduced, virtually non-detectable DCK levels (Figure 6A). Among twelve single cell-derived clones of HL60 and PL21, none displayed similarly low DCK levels (Figure 6A).

**Figure 6.**
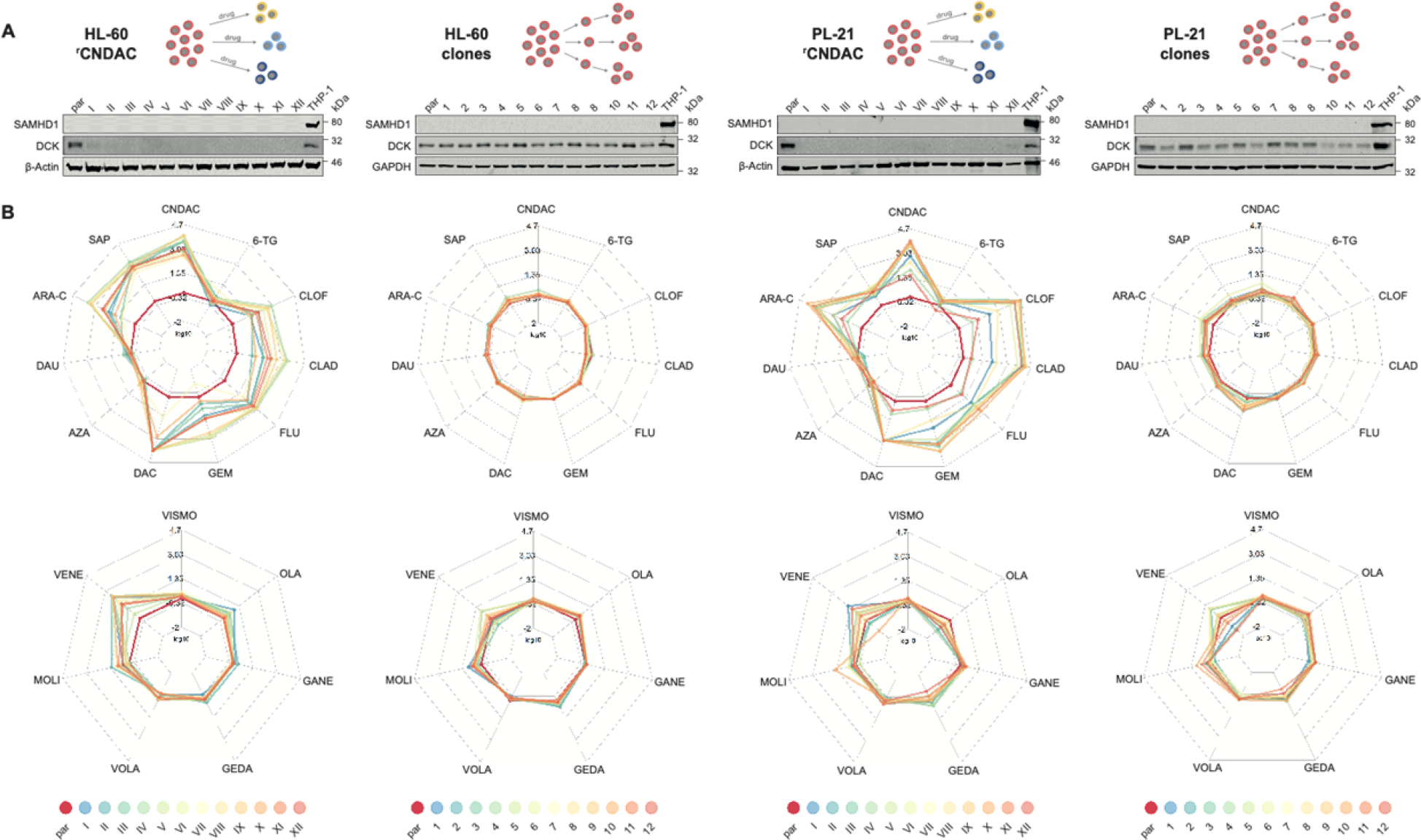
Acquired resistance to CNDAC is associated with decreased DCK levels and accompanied by cross-resistance to DCK-dependent nucleoside analogues. (A) Schematic illustrations of the establishment of CNDAC-resistant HL-60 and PL-21 cells by step-wise increasing drug concentrations during cell culture and of the establishment of single cell-derived clones by limited dilution. Moreover, representative Western blots indicating SAMHD1 and DCK levels in CNDAC-adapted HL-60 (HL-60^r^CNDACI-XII) and PL21 (PL21^r^CNDACI-XII) sublines and in single cell-derived clonal sublines of these cell lines. GAPDH and β-Actin served as loading controls. (B) Resistance profiles of CNDAC-adapted HL-60 and PL-21 sublines and single cell-derived clones of HL-60 and PL-21. Upper spider webs show sensitivity to the cytotoxic drugs CNDAC, 6-Thioguanine (6-TG), Clofarabine (CLOF), Cladribine (CLAD), Fludarabine (FLU), Gemcitabine (GEM), Decitabine (DAC), 5-Azacytidine (AZA), Daunorubicin (DAU), Cytarabine (ARA-C), and Sapacitabine (SAP), while lower spider webs display sensitivity to the targeted drugs Vismodegib (VISMO), Olaparib (OLA), Ganetespib (GANE), Gedatolisib (GEDA), Volasertib (VOLA), Molibresib (MOLI), and Venetoclax (VENE). Values are depicted as fold changes in drug concentrations that reduce cell viability by 50% (IC_50_s) between the respective parental AML cell line (shown in red) and the resistant cell lines or clones. Points closer to the centre than red lines indicate higher sensitivity to drugs in CNDAC-resistant sublines or clonal sublines than in parental cell lines, while points lying outside red lines indicate reduced sensitivity to the respective drug. IC_50_ fold changes are shown as means from three independent experiments. Numerical values are provided in Supplementary Table 6.

Then, we determined resistance profiles in the CNDAC-resistant HL60 and PL21 sublines and the clonal HL60 and PL21 sublines to a set of cytotoxic (CNDAC, sapacitabine, cytarabine, clofarabine, cladribine, fludarabine, gemcitabine, decitabine, azacytidine, 6-thioguanine, daunorubicin) and targeted (venetoclax, vismodegib, olaparib, ganetespib, volasertib, gedatolisib, molibresib) drugs (Figure 6B, Suppl. Table 6).

In addition to resistance to CNDAC and its prodrug sapacitabine, all CNDAC-adapted sublines also consistently displayed a markedly reduced sensitivity to the nucleoside analogues clofarabine, cladribine, fludarabine, gemcitabine, and decitabine, whose activation critically depends on monophosphorylation by DCK (Figure 6B, Suppl. Table 6). In contrast, there was no cross-resistance to the nucleoside analogues azacytidine and 6-thioguanine that are no DCK substrates and to the anthracycline daunorubicin. This suggests that that reduced *DCK* expression is the predominant acquired resistance mechanism in our panel of CNDAC-adapted AML cell lines.

This notion was also confirmed by the general lack of cross-resistance to targeted drugs with a range of different targets, including the smoothend receptor (vismodegib), PARP1 (olaparib), HSP90 (ganetespib), PLK1 (volasertib), and PI3K/mTOR (gedatolisib). There was some level of resistance to the BET inhibitor molibresib among the CNDAC-adapted sublines (Figure 6B, Suppl. Table 6). However, some level of resistance to these drugs was also detected among the clonal HL60 and PL21 sublines (Figure 6B, Suppl. Table 6), which may suggest that this molibresib resistance may rather be the consequence of clonal selection processes during resistance formation and not part of the acquired CNDAC resistance mechanisms.

The Bcl-2 inhibitor venetoclax was the only targeted drug against which the CNDAC-adapted sublines displayed an increased level of resistance that was not detectable in the clonal sublines (Figure 6B, Suppl. Table 6). This may indicate a generally increased resistance to apoptosis in the CNDAC-adapted sublines (Figure 6B, Suppl. Table 6), which may reflect that apoptosis induction is anticipated to be part of the anti-cancer mechanism of action of CNDAC [Liu et al., 2019].

Taken together, our findings suggest that DCK downregulation is the major acquired CNDAC resistance mechanism in AML cells, potentially complemented by a generally reduced potential to undergo apoptosis.

### Role of SAMHD1 and DCK in CNDAC cross-resistance of AML cell lines adapted to drugs from different classes

In contrast to the CNDAC-adapted AML cell lines introduced here, which displayed reduced *DCK* expression as main acquired resistance mechanism, AML cell lines adapted to the SAMHD1 substrates cytarabine or decitabine were characterised by a combination of increased SAMHD1 levels and decreased DCK levels [Schneider et al., 2017; Oellerich et al., 2019].

CNDAC-adapted AML sublines displayed pronounced cross-resistance to nucleoside analogues that are activated by DCK but not to anti-leukaemia drugs with other mechanisms of action (Figure 6). In a reversed setting, we next investigated CNDAC in a panel consisting of the AML cell line HL60 and its sublines adapted to the nucleoside analogues cytarabine, araG, azacytidine, and fludarabine, the purine antagonist 6-mercaptopurine, the Bcl-2 inhibitor venetoclax, the PARP inhibitor olaparib, and the polo-like kinase 1 inhibitor volasertib.

The nucleoside analogue-resistant HL60 sublines displayed increased SAMHD1 and/ or decreased DCK levels (Figure 7A) and pronounced CNDAC resistance (Figure 7B, Suppl. Figure 9), while little or no CNDAC resistance was detected in the remaining sublines (Figure 7A, Figure 7B, Suppl. Figure 9). Moreover, cellular SAMHD1 levels directly and cellular DCK levels inversely correlated with the CNDAC IC_50_s (Figure 7C), indicating that enhanced SAMHD1 levels and reduced DCK levels contribute to cross-resistance to CNDAC. VPX-VLP-mediated SAMHD1 depletion sensitised nucleoside analogue-adapted HL60 sublines to CNDAC to various extents (Figure 7D), which probably reflects the relative importance of SAMHD1 and DCK levels for CNDAC resistance in these cell lines.

**Figure 7.**
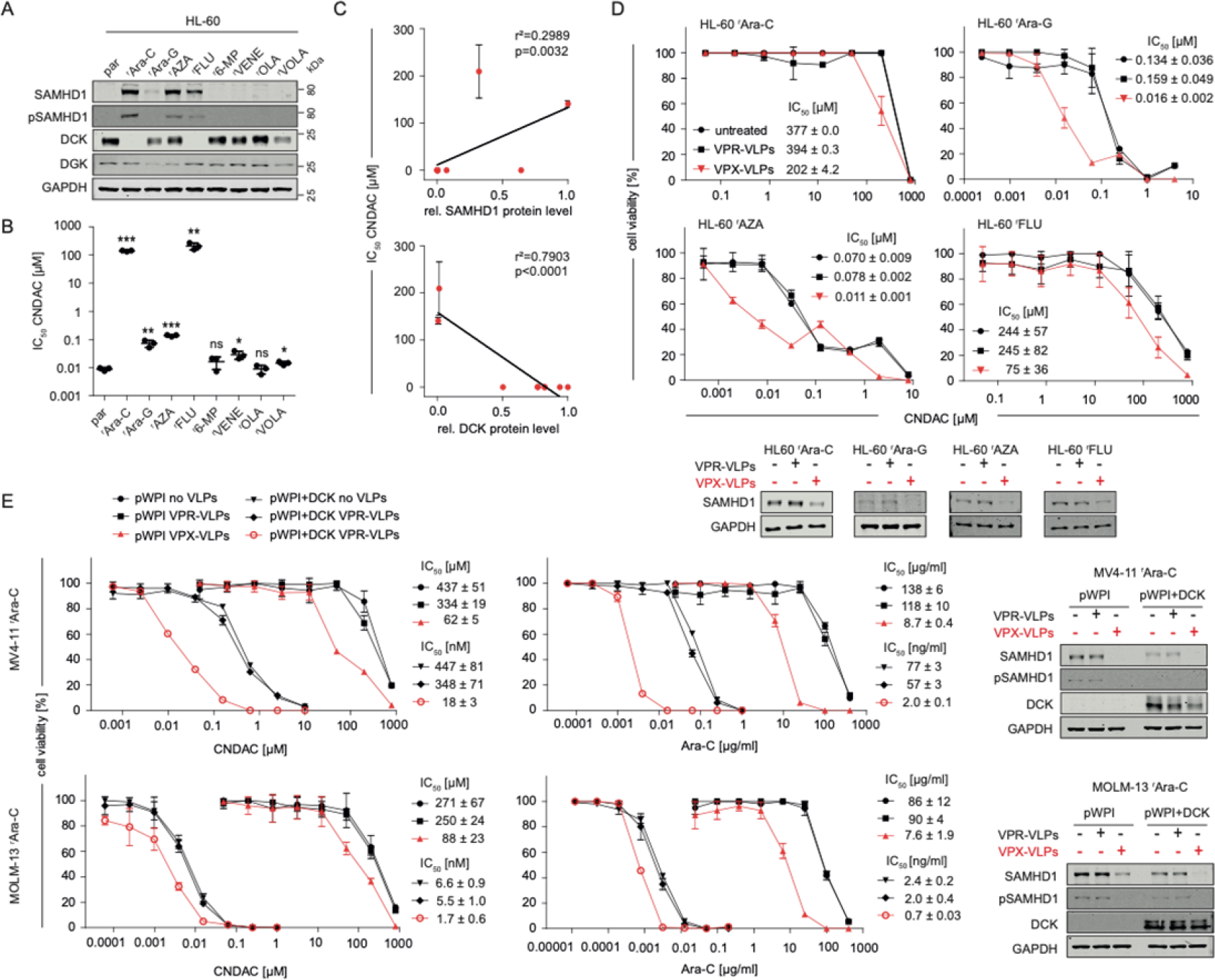
SAMHD1 and DCK regulate CNDAC cross-resistance of AML cell lines adapted to drugs from different classes. (A) Representative Western blots of SAMHD1, phosphorylated SAMHD1 (pSAMH1), DGK, and DCK in HL-60 sublines adapted to cytarabine (Ara-C), arabinosylguanine (Ara-G), 5-azacytidine (AZA), fludarabine (FLU), 6-mercaptopurine (6-MP), venetoclax (VENE), olaparib (OLA), and volasertib (VOLA). GAPDH served as loading control. (B) CNDAC concentrations that reduce cell viability by 50% (IC_50_s) in drug-adapted HL-60 sublines. Horizontal lines and error bars represent means ± SD of three independent experiments, each performed in three technical replicates. p-values were determined by two-tailed Student’s t-tests (*p < 0.05; **p < 0.01; ***p < 0.001). (C) Correlation of the CNDAC IC_50_ values with cellular SAMHD1 or DCK protein levels, quantified using the near-infrared Western blot image shown in (A) to determine the ratio SAMHD1/ GAPDH or DCK/ GAPDH. (D) CNDAC dose-response curves in drug-adapted HL-60 sublines in the absence or presence of VPX virus-like particles (VPX-VLPs, cause SAMHD1 depletion) or VPR virus-like particles (VPR-VLPs, negative control). Each symbol represents the mean ± SD of three technical replicates of one representative experiment out of three. Concentrations that reduce AML cell viability by 50% (IC_50_s) ± SD and Western Blots showing SAMHD1 degradation by VPX-VLPs are provided. (E) CNDAC or cytarabine (Ara-C) dose-response curve in cytarabine-adapted MV4-11 or MOLM-13 cells (characterised by loss of *DCK* expression) stably transduced with either DCK (pWpi+DCK) or an empty vector (pWPI) in the absence or presence of VPX virus-like particles (VPX-VLPs), or VPR virus-like particles (VPR-VLPs). Each symbol represents the mean ± SD of three technical replicates of one representative experiment out of three. IC_50_s (mean ± SD) and Western Blots showing successful transduction with DCK and SAMHD1 degradation by VPX-VLPs are provided.

Next, we used cytarabine-adapted MV4-11 and MOLM13 sublines to further study the role of SAMHD1 and DCK in cross-resistance of nucleoside analogue-adapted AML cells to CNDAC (Figure 7E). In both cell lines, VPX-VLP-mediated SAMHD1 depletion resulted in reduced CNDAC IC_50_s, which further decreased upon forced *DCK* expression. Similar findings were obtained with regard to the cytarabine resistance in these two cell lines (Figure 7E). This confirms that, in principle, cellular SAMHD1 and DCK levels are involved in determining AML cell sensitivity to CNDAC (and cytarabine), although, as shown in this study, intrinsic and acquired CNDAC resistance differ in AML cells in that intrinsic CNDAC resistance is predominantly driven by high SAMHD1 levels and acquired CNDAC resistance by a reduction in DCK.

## Discussion

The findings of this study indicate that in AML cells intrinsic CNDAC resistance is predominantly driven by SAMHD1, whereas acquired CNDAC resistance is primarily caused by reduced DCK levels. This difference is of potential clinical significance, because SAMHD1 is a candidate biomarker for predicting CNDAC sensitivity in therapy-naïve patients, while DCK is a candidate biomarker for the early detection of resistance formation.

SAMHD1 is known to interfere with the activity of a range of anti-cancer nucleoside analogues as hydroxylase that cleaves the activated nucleoside analogue triphosphates [Schneider et al., 2017; Herold et al., 2017; Knecht et al., 2018; Oellerich et al., 2019; Rothenburger et al., 2020; Xagorias et al., 2021]. The finding that SAMHD1 levels critically determine AML (and ALL) cell sensitivity to CNDAC is nevertheless somewhat unexpected, as CNDAC had originally been suggested to be a SAMHD1 inhibitor [Hollenbaugh et al., 2017].

However, data from a large range of cell line models (including clonal AML sublines characterised by varying SAMHD1 levels) and patient samples demonstrated that high SAMHD1 levels are associated with reduced CNDAC sensitivity and that CRISPR/Cas9-, siRNA-, and VPX-VPL (virus-like particles carrying the lentiviral VPX protein)-mediated SAMHD1 depletion increase cellular CNDAC-TP levels and sensitise AML cells to CNDAC. In agreement, enzymatic assays and crystallisation studies showed that CNDAC-TP is cleaved by SAMHD1, but can in contrast to some other nucleoside analogues [Ji et al., 2013; Ji et al., 2014; Hollenbaugh et al., 2017; Knecht et al., 2018; Oellerich et al., 2019] not activate SAMHD1 via binding to the A2 site.

Moreover, the determination of physiological dNTPs in the presence of CNDAC and combination experiments with the SAMHD1 substrate cytarabine did not provide evidence that CNDAC may function as pharmacological SAMHD1 inhibitor in leukaemia cells.

Although cellular SAMHD1 levels, but not those of DCK that is critical for CNDAC phosphorylation and activation [Lotfi et al., 2003; Homminga et al., 2011; Wu et al., 2021], predominantly determined CNDAC sensitivity in CNDAC-naïve cells, the establishment of 24 CNDAC-resistant AML sublines unanimously resulted in a loss of DCK but not in an increase of SAMHD1. This differs from acquired resistance mechanisms against the nucleoside analogues cytarabine and decitabine that were found to include both increased SAMHD1 levels and decreased DCK levels [Schneider et al., 2017; Oellerich et al., 2019]. Two previously established CNDAC-adapted cancer cell lines had been shown to display reduced DCK levels but a contribution of SAMHD1 had not been investigated [Obata et al., 1998; Obata et al., 2001].

CNDAC-adapted AML sublines consistently displayed cross-resistance to other nucleoside analogues known to be activated by DCK but no pronounced cross-resistance to other drugs with various mechanisms of action, further indicating that loss of DCK is the crucial resistance mechanism in CNDAC-adapted cells. Moreover, these data also show that drugs, which do not depend on DCK for activation, remain viable treatment options after resistance acquisition to CNDAC.

Similarly, among AML sublines adapted to a range of different anti-cancer drugs, only nucleoside analogues that displayed increased SAMHD1 and/ or decreased DCK levels were less sensitive to CNDAC. Thus, acquired resistance to a range of different anti-leukaemic drugs is unlikely to affect the efficacy of CNDAC.

Cytarabine- and decitabine-adapted AML cell lines are characterised by a combination of increased SAMHD1 levels and/ or reduced DCK levels as demonstrated previously [Schneider et al., 2017; Oellerich et al., 2019]. Although acquired CNDAC resistance was mediated by decreased DCK levels, both increased SAMHD1 levels and decreased DCK levels contributed to cross-resistance of cytarabine-adapted cells to CNDAC. In the future, it will be interesting to investigate why acquired resistance mechanisms differ between CNDAC-adapted cells on the one hand and cytarabine- and decitabine-adapted cells on the other hand.

In conclusion, intrinsic AML cell response to CNDAC critically depends on cellular SAMHD1 levels, whereas acquired CNDAC resistance is predominantly mediated by reduced DCK levels. This adds to data demonstrating differences between intrinsic and acquired resistance mechanisms [Michaelis et al., 2019; Oellerich et al., 2019; Santoni-Rugiu et al., 2019; Touat et al., 2020]. These findings also indicate that SAMHD1 is a candidate biomarker predicting CNDAC response in the intrinsic resistance setting, while DCK plays a potential role as biomarker indicating therapy failure early in the acquired resistance setting. Moreover, CNDAC-adapted cells displayed no or limited cross-resistance to drugs whose activity is not influenced by DCK or SAMHD1. Similarly, CNDAC was still effective in cells adapted to drugs that are not affected by DCK or SAMHD1. These findings indicate treatment options after therapy failure.

## Methods

### Compounds

CNDAC was purchased from biorbyt (via Biozol, Eching, Germany), 5-azacytidine, cytarabine, cladribine, clofarabine, decitabine, and fludarabine from Tocris Biosciences (via Bio-Techne GmbH, Wiesbaden, Germany), 6-thioguanine, ganetespib, molibresib, olaparib, sapacitabine, venetoclax, and vismodegib from MedChemExpress (via Hycultec, Beutelsbach, Germany), daunorubicin, gedatolisib, and volasertib from Selleckchem (Berlin, Germany), gemcitabine from Hexal (Holzkirchen, Germany), GTP and dATP from Thermo Scientific (Dreieich, Germany), and CNDAC-TP from Jena Bioscience GmbH (Jena, Germany).

### Cell culture

The human AML cell lines HEL (DSMZ No. ACC 11), HL-60 (DSMZ No. ACC 3), KG-1 (DSMZ No. ACC 14), ML-2 (DSMZ No. ACC 15), MOLM-13 (DSMZ No. ACC 554), MONO-MAC-6 (DSMZ No. ACC 124), MV4-11 (DSMZ No. ACC 102), NB-4 (DSMZ No. ACC 207), OCI-AML-2 (DSMZ No. ACC 99), OCI-AML-3 (DSMZ No. ACC 582), PL-21 (DSMZ No. ACC 536), SIG-M5 (DSMZ No. ACC 468), and THP-1 (DSMZ No. ACC16) and the human ALL cell lines 697 (DSMZ No. ACC 42), ALL-SIL (DSMZ No. ACC 511), BALL-1 (DSMZ No. ACC 742), CTV-1 (DSMZ No. ACC 40), GRANTA-452 (DSMZ No. ACC 713), HAL-01 (DSMZ No. ACC 610), HSB-2 (DSMZ No. ACC 435), JURKAT (DSMZ No. ACC 282), KE-37 (DSMZ No. ACC 46), MHH-CALL-4 (DSMZ No. ACC 337), MN-60 (DSMZ No. ACC 138), MOLT-4 (DSMZ No. ACC 362), MOLT-16 (DSMZ No. ACC 29), NALM-6 (DSMZ No. ACC 128), NALM-16 (DSMZ No. ACC 680), P12-ICHIKAWA (DSMZ No. ACC 34), REH (DSMZ No. ACC 22), ROS-50 (DSMZ No. ACC 557), RPMI-8402 (DSMZ No. ACC 290), RS4;11 (DSMZ No. ACC 508), SEM (DSMZ No. ACC 546), TANOUE (DSMZ No. ACC 399), and TOM-1 (DSMZ No. ACC 578) were obtained from DSMZ (Deutsche Sammlung von Mikroorganismen und Zellkulturen GmbH, Braunschweig, Germany). The ALL cell line CCRF-CEM (ATCC No. CCL-119) was received from ATCC (Manassas, VA, US), the ALL cell line KARPAS231 from Cambridge Enterprise Ltd. (Cambridge, UK), and the ALL cell line J-JHAN was kindly provided by Professor R. Tedder (University College London) [Cinatl et al., 1995].

Drug-resistant cell sublines were established by continuous exposure of sensitive parental cell lines HL-60 and PL-21 to step-wise increasing drug concentrations, as previously described [Michaelis et al., 2011] and are part of the Resistant Cancer Cell Line (RCCL) collection (https://www.kent.ac.uk/stms/cmp/RCCL/RCCLabout.html) [Michaelis et al., 2019]. Briefly, cells were cultured at increasing drug concentrations, starting with concentrations that inhibited the viability of the parental cell lines by 50% (IC_50_). Drug concentrations were increased every 2 to 6 weeks until cells readily grew in the presence of the drug. In this way 12 independent CNDAC-resistant sublines of HL-60 and PL-21 were generated each and designated as HL-60^r^CNDAC^200nM^ I – XII and PL-21 ^r^CNDAC^2µM^ I-XII. HL-60 cells with acquired resistance to the drugs Cytarabine (Ara-C), arabinosylguanine (AraG), 5-azacytidine (AZA), fludarabine (FLUDA), 6-Mercaptopurine (6-MP), venetoclax (VENE), olaparib (OLA), and volasertib (VOLA) were designated as HL-60^r^AraC^2µg/ml^, HL-60^r^AraG^100µM^, HL-60^r^5-AZA^1µM^, HL-60^r^FLUDA^1µg/ml^, HL-60^r^6-MP^2µM^, HL-60^r^VENE^2µM^, HL-60^r^OLA^20µM^ and HL-60^r^VOLA^200nM^.

Clonal sublines were generated by limiting dilution. Cells were plated at a density of 1 cell per well on a 96-well plate and grown for 1 - 2 weeks. Wells with only one visible cell colony were identified and the respective clones were expanded.

SAMHD1-deficient THP-1 (THP-1 KO) cells and control cells (THP-1 CTRL) were generated using CRISPR/Cas9 approach as previously described [Wittmann et al., 2015; Schneider et al., 2017; Oellerich et al., 2019]. THP-1 cells were plated at a density of 2 × 10^5^ cells/ mL. After 24h, 2.5 × 10^6^ cells were suspended in 250µl Opti-MEM, mixed with 5µg CRISPR/Cas plasmid DNA, and electroporated in a 4-mm cuvette using an exponential pulse at 250 V and 950 mF in a Gene Pulser electroporation device (Bio-Rad Laboratories, Feldkirchen, Germany). We used a plasmid encoding a CMV-mCherry-Cas9 expression cassette and a human SAMHD1 gene specific gRNA driven by the U6 promoter. An early coding exon of the SAMHD1 gene was targeted using the following gRNA construct: 5′-CGGAAGGGGTGTTTGAGGGG-3′. Cells were allowed to recover for 2 days in 6-well plates filled with 4 ml medium per well before being FACS sorted for mCherry-expression on a BD FACS Aria III (BD Biosciences, Heidelberg, Germany). For subsequent limiting dilution cloning, cells were plated at a density of 5, 10, or 20 cells per well of nine round-bottom 96-well plates and grown for 2 weeks. Plates were scanned for absorption at 600 nm and growing clones were identified using custom software and picked and duplicated by a Biomek FXp (Beckman Coulter, Krefeld, Germany) liquid handling system.

DCK-expressing MV4-11rAraC^2µg/ml^ and MOLM-13rAraC^2µg/ml^ cells were established by lentiviral transduction and designated as MV4-11rAraC^2µg/ml^-pWPI+DCK and MOLM-13rAraC^2µg/ml^-pWPI+DCK (or MV4-11rAraC^2µg/ml^-pWPI and MOLM-13rAraC^2µg/ml^-pWPI for control cells transduced with the empty vector). To generate the pWPI+DCK plasmid, the dCK gene was PCR-amplified from pDNR-Dual_dCK (DNAsu HsCD00000962) using Pfu DNA polymerase (Promega, Germany) and gene-specific primers (Eurofins Genomics, Germany) and subcloned into pWPI IRES puro via BamHI/SpeI. The plasmid was verified by Sanger sequencing (Eurofins Genomics, Germany). For the generation of lentiviral vectors 293T cells were co-transfected with pWPI+DCK (or pWPI as control), Addgene packaging plasmid pPAX, an envelope plasmid encoding VSV-G and pAdVAntage (Promega). Four days after transfection, lentiviral vectors were harvested and concentrated by ultracentrifugation. For lentiviral transduction MV4-11rAraC^2µg/ml^ and MOLM-13rAraC^2µg/ml^ cells were seeded at 5 x 10^5^ cells/ well of a 96-well-plate and spinoculated with the lentiviral vectors. 24 hours after transduction, successfully transduced cells were selected with 3 µg/ml puromycin (Sigma-Aldrich) and DCK expression was monitored by Western Blot.

All cell lines were cultured in IMDM (Biochrom, Cambridge, UK) supplemented with 10% FBS (SIG-M5 20% FBS, Sigma-Aldrich, Taufkirchen, Germany), 4 mM L-Glutamine (Sigma-Aldrich), 100 IU/ml penicillin (Sigma-Aldrich), and 100 mg/ml streptomycin (Sigma-Aldrich) at 37°C in a humidified 5% CO2 incubator. Cell lines were routinely tested for Mycoplasma, using the MycoAlert PLUS assay kit from Lonza (Basel, Switzerland), and were authenticated by short tandem repeat profiling.

### Primary AML samples

Peripheral blood or bone marrow samples derived from AML patients between 2018 and 2020 were obtained from the UCT Biobank of the University Hospital Frankfurt. The use of peripheral blood and bone marrow aspirates was approved by the Ethics Committee of Frankfurt University Hospital (approval no. SHN-03-2017). All patients gave informed consent to the collection of samples and to the scientific analysis of their data and of biomaterial obtained for diagnostic purposes according to the Declaration of Helsinki.

Mononuclear cell (MNC) fractions were purified by gradient centrifugation with Biocoll cell separation solution (Merck Millipore, Darmstadt, Germany). Leukemic cells were enriched by negative selection with a combination of CD3-, CD19- and CD235a-microbeads (all obtained from Miltenyi Biotec, Bergisch Gladbach, Germany, 130-050-301, 130-050-101, 130-050-501) according to the manufacturer’s instructions and separated by the autoMACS™ Pro Separator (Miltenyi Biotec). FACS staining and treatment for viability assays of AML blasts was executed immediately after isolation. Culture medium for AML blasts consisted of IMDM (Biochrom) supplemented with 10% FBS, 4 mM L-glutamine, 25 ng/ml hTPO, 50 ng/ml hSCF, 50 ng/ml hFlt3-Ligand and 20 ng/ml hIL-3 (all obtained from Miltenyi Biotec, 130-094-013, 130-096-695, 130-096-479, 130-095-069).

### Viability assay

The viability of AML and ALL cell lines treated with various drug concentrations was determined by 3-(4,5-dimethylthiazol-2-yl)-2,5-diphenyltetrazolium bromide (MTT) assay modified after Mosman [Mosmann, 1983], as previously described [Onafuye et al., 2019]. Cells suspended in 100 µL cell culture medium were plated per well in 96-well plates and incubated in the presence of various drug concentrations for 96 h. Then, 25 µL of MTT solution (2 mg/mL (w/v) in PBS) were added per well, and the plates were incubated at 37 °C for an additional 4 h. After this, the cells were lysed using 100 µL of a buffer containing 20% (w/v) sodium dodecylsulfate in 50% (v/v) N,N-dimethylformamide with the pH adjusted to 4.7 at 37 °C for 4 h. Absorbance was determined at 570 nm for each well using a 96-well multiscanner (Tecan Spark, Tecan, Crailsheim, Germany). After subtracting of the background absorption, the results are expressed as percentage viability relative to control cultures which received no drug. Drug concentrations that inhibited cell viability by 50% (IC_50_) were determined using CalcuSyn (Biosoft, Cambridge, UK) or GraphPad Prism (San Diego, CA, USA).

For AML blasts viability assays were performed using the CellTiter-Glo (Promega, Walldorf, Germany) assay according to the manufacturer’s protocol. Briefly, cells were seeded at 5,000 cells per well in 96-well plates and treated for 96 hours. Luminescence was measured on a Tecan Spark (Tecan). IC_50_ values were calculated usind GraphPad Prism.

### Caspase 3/7 assay

To determine Caspase 3/7 activity in THP-1 SAMHD1 KO and CTRL cells the Caspase-Glo 3/7 assay (Promega, Walldorf, Germany) was used according to the manufacturer’s protocol. Briefly, cells were seeded at 5,000 cells per well in white 96-well plates, treated with different concentrations of CNDAC and incubated for 24, 48 and 72 hours at 37°C in a humidified 5% CO_2_ incubator. After incubation an equal volume of Caspase-Glo 3/7 reagent was added, mixed for 30 minutes and luminescence was measured on a Tecan Spark (Tecan).

### Determination of Population doubling time (PDT)

To generate a growth curve, cells were seeded at 2,000 cells per well in a white 96-well plate in 100µl culture medium and incubated for 0, 1, 2, 3, 4 and 7 days at 37°C in a humidified 5% CO_2_ incubator. Cell viability was detected using the CellTiter-Glo assay (Promega) according to the manufacturer’s protocol. Growth curves were created and the population doubling times calculated using the following formula:

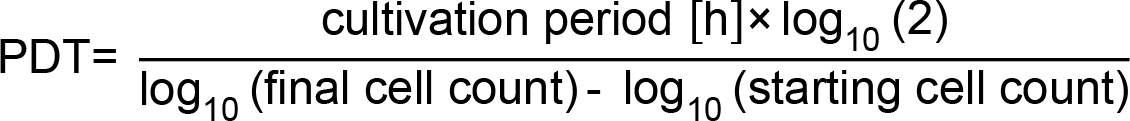

### Western blot analysis

Whole-cell lysates were prepared by using Triton-X sample buffer containing protease inhibitor cocktail from Roche (Grenzach-Wyhlen, Germany). The protein concentration was assessed by using the DC Protein assay reagent obtained from Bio-Rad Laboratories. Equal protein loads were separated by sodium dodecyl sulfate-polyacrylamide gel electrophoresis and proteins were transferred to nitrocellulose membranes (Thermo Scientific, Dreieich, Germany). The following primary antibodies were used at the indicated dilutions: SAMHD1 (Proteintech, St. Leon-Roth, Germamy, 12586-1-AP, 1:1000), β-actin (BioVision, Milpitas, CA, US, 3598R-100, 1:5000), pSAMHD1 (Cell Signaling, Frankfurt am Main, Germany, 89930S, 1:1000), and GAPDH (Trevigen via Bio-Techne, Wiesbaden, Germany, 2275-PC-10C, 1:5000), DCK (abcam, Berlin, Germany, ab96599, 1:4000), DGK (Santa Cruz Biotechnology, Heidelberg, Germany, sc-398093, 1:100), PARP (Cell Signaling, 9542S, 1:1000), H2AX (Cell Signaling, 2595S, 1:1000), γH2AX (Cell Signaling, 9718S, 1:1000), Chk2 (Cell Signaling, 2662S, 1:1000), pChk2 (Cell Signaling, 2661S, 1:1000), TIF-1β (Cell Signaling, 4124S, 1:1000), pTIF-1β (Cell Signaling, 4127S, 1:1000). Visualisation and quantification were performed using IRDye-labeled secondary antibodies (LI-COR Biotechnology, Bad Homburg, Germany, IRDye®800CW Goat anti-Rabbit, 926-32211 and IRDye®800CW Goat anti-Mouse IgG, 926-32210) according to the manufacturer’s instructions. Band volume analysis was conducted by Odyssey LICOR.

### Flow Cytometry

The intracellular SAMHD1 staining of AML blasts was performed as previously described [Baldauf et al., 2012] with SAMHD1-antibody from Proteintech (12586-1-AP, 1:100). Staining for surface markers (CD33, CD34, CD45) for AML blasts was applied before fixation with the following fluorochrome-conjugated antibodies: CD33-PE and CD34-FITC, both from Miltenyi Biotech (130-111-019, 130-113-178) and CD45-V450 from BD Pharmingen (Heidelberg, Germany, 642275), all diluted 1:5 per 1 x 10^7^ cells, and goat anti-rabbit Alexa-Fluor-660 from Invitrogen, Life technologies (1:200, A-21073) as secondary antibody for SAMHD1 staining. Samples were analysed by using a FACSVerse flow cytometer from BD Biosciences (Heidelberg, Germany) and the FlowJo software (FlowJo LLC, Ashland, OR, US). To determine the mean fluorescence intensity (MFI) for SAMHD1, the geometric mean for the isotype control was subtracted from the geometric mean for SAMHD1.

### SAMHD1 promoter methylation

The *SAMHD1* promoter contains five *HpaII* sites surrounding the transcription start site [de Silva et al., 2013]. To measure methylation of the *SAMHD1* promoter genomic DNA was treated with the methylation-sensitive HpaII endonuclease as described previously [de Silva et al., 2013; Oellerich et al., 2019]. Methylation of the *HpaII* sites in the *SAMHD1* promoter prevents digestion by HpaII and the intact sequence serves then as a template for PCR amplification using *SAMHD1* promoter-specific primers that flank the *HpaII* sites: PM3.fwd: TTCCGCCTCATTCGTCCTTG and PM3.rev: GGTTCTCGGGCTGTCATCG were used as SAMHD1 promoter-specific primers. A single PCR product (993-bp) corresponding to the *SAMHD1* promoter sequence was obtained from untreated genomic DNA and treated DNA from cells with methylated but not from cells with unmethylated *SAMHD1* promoter. To serve as input control, a 0.25-kb fragment of the *GAPDH* gene lacking *HpaII* sites was PCR-amplified using the same template DNA.

### Manipulation of cellular SAMHD1 levels using siRNA or Vpx-VLPs

For siRNA-mediated silencing, AML blasts (1 x 10^6^) were transfected with 2.5 µM ON-TARGET plus human SAMHD1 siRNA SMART-pool obtained from Dharmacon (Munich, Germany, L-013950-01-0050) in resuspension electroporation buffer R (Invitrogen, Dreieich, Germany) using the Neon transfection system (Invitrogen) according to the manufacturer’s recommendation. Additionally, ON-TARGET plus Non-targeting Control Pool obtained from Dharmacon (D-001810-10-50) was transfected in parallel. The electroporation was performed with one 20 msec pulse of 1700 V and analysed 48 h after transfection by western blotting and a cell viability assay.

For Vpx virus-like particle (VLP)-mediated SAMHD1 degradation, cells were spinoculated with VSV-G pseudotyped virus-like particles carrying either Vpx or Vpr as control from SIVmac251. VLPs carrying Vpx or Vpr were produced by co-transfection of 293T cells with pSIV3 + *gag pol* expression plasmids and a plasmid encoding VSV-G as previously described [Schneider et al., 2017; Oellerich et al., 2019]. For viability assays cells were preincubated with VLPs for 24 h before the studied compounds were added.

### LC-MS/MS Analysis

AML or ALL cells were seeded at 2,5 x 10^5^ cells per well in 24 well plates, treated with 10µM CNDAC and incubated at 37°C in a humidified 5% CO2 incubator for 6 h. Subsequently, cells were washed twice in 1 ml PBS, pelleted and stored at - 80°C until measurement. The concentrations of canonical dNTPs and CNDAC-triphosphate in the samples were analysed by liquid chromatography-electrospray ionization-tandem mass spectrometry, as previously described for canonical dNTPs [Thomas et al., 2015]. Briefly, the analytes were extracted by protein precipitation with methanol. An anion-exchange HPLC column (BioBasic AX, 150 x 2.1 mm, 5 µM, Thermo Scientific) was used for the chromatographic separation and a 5500 QTrap (Sciex, Darmstadt, Germany) was used as analyser, operating as triple quadrupole in positive multiple reaction monitoring (MRM) mode. CNDAC-TP was quantified using 2-deoxycytidine-^13^C_9_,^15^N_3_-triphosphate (^13^C_9_,^15^N_3_-dCTP) as internal standard (IS). The precursor-to-product ion transition used as quantifier was m/z 493.1 → 112.1 for CNDAC-TP. Owing to the lack of commercially available standards for CNDAC-TP, relative quantification was performed by comparing the peak area ratios (analyte/IS) of the differently treated samples.

### Protein Expression and Purification

N-terminal 6×His-tagged SAMHD1 (residues 113 to 626, H206R D207N) was expressed in BL21 (DE3) Escherichia coli grown in Terrific Broth medium at 200 rpm, 18° C for 16 hr. Cells were re-suspended in buffer and passed through a microfluidizer. Cleared lysates were purified using nickel-nitrilotriacetic acid (Ni-NTA) affinity and size-exclusion chromatography. Proteins were stored in a buffer containing 50 mM Tris-HCl, pH 8, 150 mM NaCl, 0.5 mM TCEP, 5 mM MgCl2, and 10% glycerol.

### Crystallization and Data Collection

Purified SAMHD1 protein in buffer (50 mM Tris⋅HCl, pH 8.0, 150 mM NaCl, 5 mM MgCl2, and 0.5 mM TCEP) was mixed with 1 mM GTP, 0.1 mM dATP, and 10 mM CNDAC. All crystals were grown at 25 °C using the microbatch under-oil method by mixing 1 μL of protein (3 mg/mL) with 1 μL of crystallization buffer (100 mM succinate–phosphate–glycine (SPG) buffer, pH 7.4, 25% PEG 1500; Qiagen). Crystals were improved by streak seeding. Crystals were cryoprotected in paratone oil and frozen in liquid nitrogen. Diffraction data were collected at Advanced Photon Source beamline 24-ID-E. The data statistics are summarized in Table 1.

### Structure Determination and Refinement

Using the previously published SAMHD1 tetramer structure (PDB ID code 4BZB), with the bound nucleotides removed, as the search model, the structure was solved by molecular replacement using PHASER [Vagin & Teplyakov, 2000; McCoy et al., 2007; Winn et al., 2011]. The model was refined with iterative rounds of restrained refinement using Refmac5 [Murshudov et al., 1997], followed by rebuilding the model to the 2Fo-Fc and the Fo-Fc maps using Coot [Emsley et al., 2010]. Refinement statistics are summarised in Suppl. Table 5. Coordinates and structure factors have been deposited in the Protein Data Bank, with accession codes listed in Suppl. Table 5.

### Enzymatic assay

In vitro SAMHD1 activity was measured as described [Seamon & Stivers, 2015]. Briefly, 1µM his-tagged human SAMHD1 and 1.5µM PPase from E.coli were incubated at room temperature in 20µL reaction buffer (50mM Tris, 150mM NaCl, 1.25mM MgCl_2_, 0.5mM TCEP, 0.05% Brij-35) and different concentrations of GTP, dGTP and CNDAC-TP in a clear 384-well plate (Corning, 3700, New York, USA). Reactions were stopped by addition of 20µL EDTA (20mM in water). Subsequently, 10µL malachite green reagent (Sigma-Aldrich, MAK307, Missouri, USA) were added. Absorbance was recorded at 620nm after incubating the samples for 60min at room temperature. For normalization, background subtraction of controls containing the same substrate and PPase concentrations but no SAMHD1 was performed.

### Statistics

Statistical data analysis was performed using GraphPad Prism. Pearson’s correlation coefficient was used to compute correlations between variables, using a t-test to assess significance of the correlation. Group comparisons were performed using Student’s t-test.

## Declarations

### Ethics approval and consent to participate

The use of peripheral blood and bone marrow aspirates was approved by the Ethics Committee of Frankfurt University Hospital (approval no. SHN-03-2017). All patients gave informed consent to the collection of samples and to the scientific analysis of their data and of biomaterial obtained for diagnostic purposes according to the Declaration of Helsinki.

## Consent for publication

Not applicable.

## Availability of data and materials

The atomic coordinates and structure factors have been deposited in the Protein Data Bank, www.wwpdb.org. The PDB ID code will be added upon publication. The Preliminary Full wwPDB X-ray Structure Validation Report is provided as supplement.

## Authors’ contributions

TR, DT, YS, PRW, TP, KK, KD, JT, CS, HB, KM, FR, BB, SF, DB, RC, NF, MNW, and JC performed experiments. All authors analysed data. JC and MM conceptualised and directed the study. TR, JC, and MM wrote the initial manuscript draft. All authors read and approved the final manuscript.

## Acknowledgements

Not applicable.

